# Discovery, Interruption, and Updating of Auditory Regularities in Memory: Evidence from Low-Frequency Brain Dynamics in Human MEG

**DOI:** 10.1101/2025.03.28.645906

**Authors:** Roberta Bianco, Kaho Magami, Marcus Pearce, Maria Chait

## Abstract

During passive listening, the brain maintains a hierarchy of predictive models to monitor the statistics of its surroundings. The automatic discovery of regular patterns has been associated with a gradual increase in sustained tonic M/EEG activity, sourced in auditory, hippocampal, and frontal areas — reflecting evidence accumulation and establishment of a regularity model. Conversely, when a regular pattern is interrupted, the sustained activity drops — indicating disengagement from the model. However, how such models are established in and retrieved from memory as well as the conditions under which they are activated and interrupted remain underexplored. In this MEG experiment (N=26; both sexes), we examined how neural responses related to model ‘establishment’ and ‘interruption’ are influenced by (1) the rate of stimulus presentation (tone presentation rate 20 Hz vs. 40 Hz), and (2) the novelty of the experienced acoustic structure (novel vs resumed REG pattern). The results show that (1) the dynamics of model interruption and establishment are independent of stimulus presentation rate, and that (2) model establishment occurred much faster when an experienced vs novel pattern was presented after pattern interruption, suggesting re-activation of the stored original model facilitated by the hippocampus. (3) Finally, sustained response rises in response to pattern establishment and interruption were localized in auditory, hippocampal, and frontal sources, supporting top-down model information flow. These results unveil the temporal dynamics and neural network underlying the brain’s construction and selection of predictive models to monitor changes in sensory statistics.

**Significance statement:** Sustained neural activity in response to statistical shifts in auditory sequences reflects how the brain integrates sensory information to build and update predictive models. The present magnetoencephalography (MEG) study aims to uncover the factors that affect this updating. Using tone-pip sequences as a controlled model system, we found that the brain tracks sequence information regardless of tempo and automatically retains a memory of previously encountered patterns, facilitating detection when they re-emerge. Source localization revealed activation in auditory, hippocampal, and frontal areas, highlighting a distributed network involved in top-down model updating and predictive processing. These findings provide new insights into the neural mechanisms of statistical learning and memory, revealing how the brain adapts perception in dynamic environments.

## Introduction

Current theories of perception and cognition propose that human ability to interact with the environment—whether by understanding, engaging with, or making decisions about the surroundings—relies on the brain’s capacity to build predictive models (Friston and Kiebel, 2009; Pezzulo and Cisek, 2016; de Lange et al., 2018). These models allow us to anticipate sensory input, optimize behavior, and efficiently process complex stimuli. Although extensive evidence across multiple domains highlights their perceptual relevance and potential neural underpinnings (Garrido et al., 2009; Winkler et al., 2009; Wacongne et al., 2011; Dehaene et al., 2015; Schröger et al., 2023), the neural mechanisms underlying model maintenance and updating remain underexplored (Heilbron and Chait, 2018; Soltani and Izquierdo, 2019; Press et al., 2020).

Here, we examine model updating within the auditory modality. Unlike visual scenes that are often stable over time, sounds require continuous monitoring of evolving sensory structure therefore providing a particularly useful framework for investigating predictive processing.

Previous research has shown that the auditory system implicitly tracks regularities in sound sequences (Barczak et al., 2018; Demarchi et al., 2019; Asokan et al., 2021; Bianco et al., 2024), integrating across multiple sound dimensions (Skerritt-Davis & Elhilali, 2021) and temporal scales (Bianco et al., 2020; Baumgarten et al., 2021; Hu et al., 2024). Barascud et al. (2016) identified a neurophysiological (MEG) correlate of sequence regularity tracking in passive listeners exposed to rapid tone-pip sequences. These were organized into either random (RAN) or deterministic regular (REG) patterns, consisting of different numbers of tones (and hence complexity). They demonstrated that: **(1)** increasingly predictable patterns elicited a gradual increase in sustained MEG power; **(2)** The initial “regularity discovery” phase involved coordinated activity across auditory cortical, frontal, and hippocampal regions; **(3)** Interrupting a REG sequence by transitioning to a RAN pattern led to a marked drop in the sustained response; **(4)** Brain response latencies closely followed the predictions of an ideal observer model (IDyOM; Pearce, 2018; Harrison et al., 2020), requiring one cycle plus ∼4 additional tones to establish pattern recognition. This suggested that neural information tracking dynamically updates based on the statistical properties of incoming sensory input. Subsequent studies have replicated these findings across the age span (Southwell et al., 2017, 2024; Herrmann and Johnsrude, 2018; Southwell and Chait, 2018; Al Jaja et al., 2020; Herrmann et al., 2021, 2023; Tóth et al., 2023; Hu et al., 2024; Zhao et al., 2025) and further suggested that changes in the sustained response may reflect precision tracking (inverse variance of the predictive distribution; Zhao et al., 2025).

Using the dynamics of the sustained response as a measure of regularity tracking, the present study asks two main questions:

### How does the brain decide to “interrupt” an existing regularity model?

Barascud et al. (2016) demonstrated that transitioning from REG to RAN patterns was associated with an abrupt drop in the sustained response, hypothesized to reflect disengagement from a top-down predictive model. Interestingly, this occurred ∼200 ms after the transition. Does this delay reflect an inherent neural processing constraint or a strategic “wait-and-see” period for integrating new evidence before interrupting the existing model? Is there an adaptive mechanism balancing stability in top-down processing with flexibility in response to environmental changes? Here, we manipulated the duration of tone pips. If the observed dynamics are driven by internal models based on informational properties of the input (Baumgarten et al., 2021), response latencies should depend on information content rather than the absolute rate of tone presentation. Conversely, if latencies remain fixed, this would suggest a predominant effect of low-level biophysical constraints (Theunissen and Miller, 1995).

Additionally, we examined whether the dynamics of model discovery are influenced by prior exposure to regularity. Specifically: **Does recent experience with a REG pattern facilitate its re-discovery when the same pattern re-emerges?** This should occur if the brain integrates longer-term contextual history as well as short-term (local) statistical information when perceiving sequence structure.

## Methods

### Participants

26 paid subjects (19 female; average age, 24.7 ± 3.09 years old) participated in the experiment. None reported a history of hearing or neurological disorders. All experimental protocols received approval from the research ethics committee of University College London, and written informed consent was acquired from each participant.

### Stimuli

Stimuli were sequences of tone-pips (gated on and off with 5-ms raised cosine ramps) with frequencies drawn from 20 values equally spaced on a logarithmic scale between 222 and 2000 Hz (12% steps). Sequences containing various transitions between REG and RAN patterns (Figure 1) were generated anew on each trial and presented to the listeners in a random order with an ISI that varied uniformly between 1.5 and 2 s.

**Figure 1.**
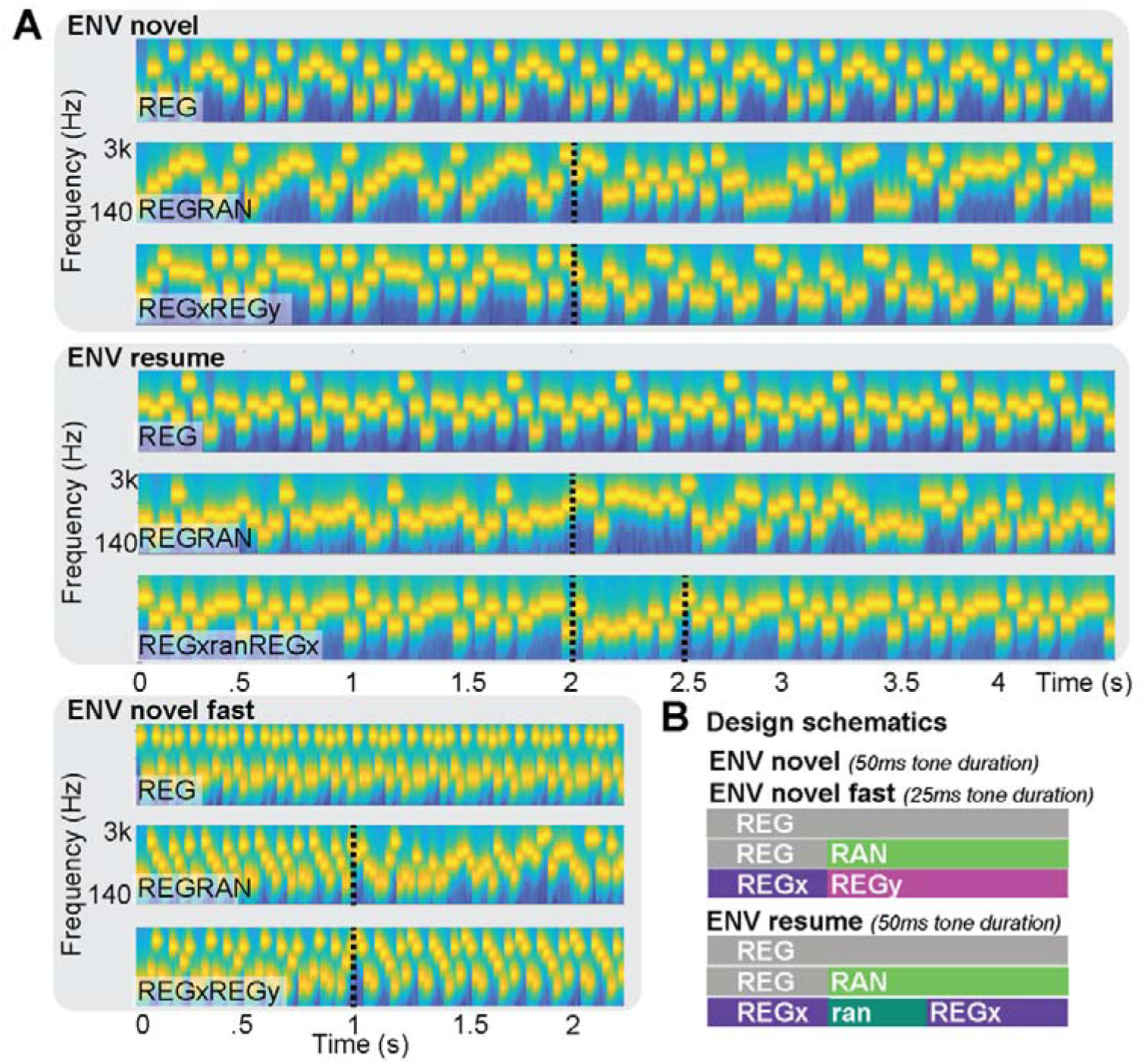
Stimuli. **A:** Three different auditory environmental contexts were presented to each participant: ENVnovel, ENVnovel fast, and ENVresume. ENVnovel included three conditions: a regular pattern (REG), a regular-to-random transition (REGRAN), and a regular-to-new regular pattern (REGxREGy). The post-transition REG (REGy) was always different from the initial REG (REGx). Sequences were composed of 50 ms tone pips. In ENVnovel fast, the same conditions were used, but with 25-ms tone-pips. In ENVresume, 50-ms tone-pips were used, with REG and REGRAN conditions as in ENVnovel, and a third condition (REGxranREGx), consisting of a regular pattern interrupted by a 0.5 s random sequence, followed by the reinstatement of the original REG pattern. **B:** Design schematics illustrating the stimuli sequences for each context and condition. Stimulus conditions within each ‘environmental context’ were equiprobable.

For each participant, we created three different sets of stimuli to simulate different auditory environmental contexts: **ENVnovel**, **ENVnovel fast**, and **ENVresume**. Each ‘context’ was blocked separately with at least a 5-minute break between contexts.

In **ENVnovel**, 50 ms tone-pips (4.5 s total sequence duration) were sequentially arranged in different orders to yield the following conditions: 1) **REG**: 10 tone-pips, randomly selected without replacement from the pool, were presented in identical cycles to form a regular sequence; 2) **REGRAN**: Containing a transition from a REG sequence (generated as described above) and a random (**RAN**) sequence generated from 10 newly sampled tones presented in random order. The transition occurred at 2 s post-onset. We ensured that the first RAN tone was always unexpected based on the preceding REG pattern and therefore constituted a sequence violation. 3) **REGxREGy**: Containing a transition (at 2 s post-onset) from a regular sequence (generated as described above) to a new regular sequence (consisting of 10 newly sampled tones but allowing for overlap with those used for REGx; as above, the first tone of REGy was always unexpected based on the preceding REGx sequence, thereby creating a sequence violation). Therefore, in this context, all transitions, REGRAN and REGxREGy, were always associated with the discontinuation of the initial REG pattern and the emergence of a new sequence. The stimuli were presented in 5 consecutive blocks. Each block consisted of 30 REG, 30 REGRAN, and 30 REGxREGy stimuli.

**ENVnovel fast**, contained the same conditions as in ENVnovel, but tones were 25 ms long (2.25 s total sequence duration). Therefore, all stimuli were presented at double the speed and were half the duration of those in ENVnovel; transitions occurred 1 s post sequence onset. The stimuli were presented in 2 consecutive blocks. Each block consisted of 50 REG, 50 REGRAN, and 50 REGxREGy stimuli.

In **ENVresume**, 50 ms tone-pips were used to form three conditions. **REG** and **REGRAN** conditions were as described in ENVnovel. **REGxranREGx** consisted of a REG sequence, presented for 2 s, then interrupted by a 0.5 s RAN sequence (10 tones chosen randomly from the pool), followed by the reinstatement of the original REG pattern. Therefore, in this context, the original REG re-occurred following the pattern interruption in 33% of the trials. The stimuli were presented in 5 consecutive blocks. Each block consisted of 30 REG, 30 REGRAN, and 30 REGxranREGx stimuli.

The order of ENVnovel and ENVresume was randomized across participants, but ENVnovel fast was consistently presented in the second position (seven participants did not perform this task because it was introduced partway through data acquisition). Sounds were stored as 16-bit .wav files at 44.1 kHz. Participants were exposed to the sounds whilst performing an incidental visual task. The task consisted of landscape images, grouped in triplets (the duration of each image was 5 s, with 2 s ISI between trials during which the screen was blank). Participants were instructed to fixate on a cross in the center of the screen and press a button whenever the third image was identical to the first image (10% trials). The visual task served as an easy decoy task for diverting subjects’ attention from the auditory stimuli. Participants were naïve to the nature of the auditory stimuli and encouraged to focus on the visual task. Feedback was displayed at the end of each block.

The experiment was controlled with the Psychophysics Toolbox for MATLAB (Kleiner et al., 2007). All auditory stimuli were presented binaurally via tube earphones (EARTONE 3A 10 Ω; Etymotic Research) inserted into the ear canal, with the volume set at a comfortable listening level, adjusted for each participant.

### Computational modeling

Stimuli were characterized using the Information Dynamics of Music (IDyOM) model (Pearce, 2018; Harrison et al., 2020), which expresses tone-by-tone unexpectedness in terms of information-theoretic surprisal. IDyOM is based on the Prediction by Partial Matching (PPM) algorithm employing a variable-order Markov model to smooth together predictions made from n-gram models of different order (context length). It generates probability distributions for each new event conditioned upon the preceding context and the prior experience of the model. Based on such distributions, the model quantifies the unexpectedness of each event as surprisal, or information content (IC) – the negative log of the conditional probability of an event. The model was run on the full stimulus set in its short-term memory configuration (STM), where the model learns incrementally based on exposure throughout each individual stimulus. The model output is summarized by averaging IC across trials for each tone position and condition, similar to how participant MEG responses are averaged.

### Data recording

Magnetic signals were recorded using CTF-275 MEG system (axial gradiometers, 274 channels; 30 reference channels; VSM MedTech). Head position was localized using three fiducial coils placed at the nasion and left/right pre-auricular points. The recording was continuous, with a sampling rate of 600 Hz and an online low-pass filter of 100 Hz. Participants were seated upright and required to remain still. The main session lasted about 1.5 hours including breaks. The stimulus presentation was divided into 12 blocks of about 5/8 min each.

### Data analysis

Analyses were conducted in MATLAB 2018b using FieldTrip (Oostenveld et al., 2011). To allow us to observe any potential effects of environmental volatility (i.e., high probability that REGx would resume following the interruption in ENVresume) on brain responses, we analyzed only the final three blocks of the ENVnovel and ENVresume conditions. In the ENVnovel fast context we analyzed the data from the two presented blocks. Data were low-pass filtered at 45 Hz (Butterworth filter, zero-phase, order 3) and segmented into epochs starting at 0.5 s pre-stimulus onset and ending at 5.5 s post-onset for ENVnovel and ENVresume and 3.75 s for ENVnovel fast. After discarding trials that were beyond twice the standard deviation from the mean (10% of trials) (NoiseTools MATLAB toolbox, de Cheveigné, 2010), denoising source separation (DSS) (de Cheveigné and Simon, 2008) was applied to maximize the reproducibility over trials.

#### Time domain analyses

For the analysis of the sequence-evoked response, data were low-pass filtered (45 Hz) but not high-pass filtered to preserve low-frequency activity. DSS was applied to all conditions together, but separately by context, (time range -0.1:4.5 s for ENVnovel and ENVresume; -0.1:2.25 s for ENVnovel fast), and the five most reproducible DSS components were projected back into sensor space.

The resulting data were summarized as root-mean-square (RMS) across 40 channels selected for each subject. The RMS is a useful summary signal, reflecting the instantaneous power of the neural response irrespective of its polarity. The mean group RMS was used for plotting.

The channels were selected based on a functional auditory localizer obtained by analyzing the M100 component of the onset response across all trials across contexts. To do so, data were epoched (-0.2:0.5 s around the trial onset), and baseline corrected to the pre-onset interval. For each subject, the 40 most strongly activated channels at the peak of the M100 (20 in each hemisphere) were considered to best reflect auditory activity (Chait et al., 2004) and selected for computing the RMS. To compare ENVnovel and ENVnovel fast, data points of the time series data were averaged into bins with each bin containing samples corresponding to the duration of a tone (Fig. 2C).

**Figure 2:**
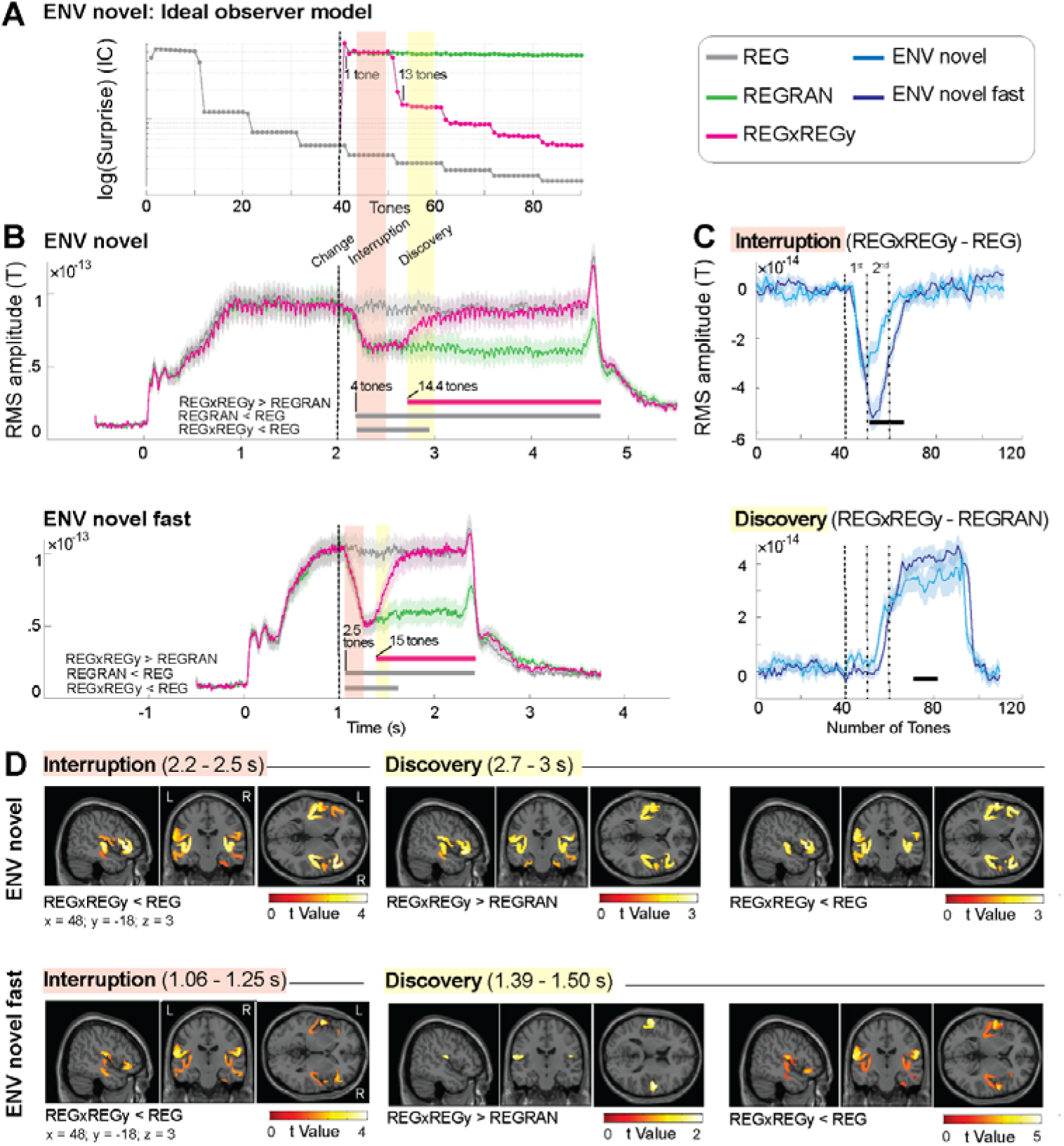
ENVnovel vs ENVnovel fast: **A:** Ideal observer model responses to all conditions in ENVnovel showing the average information content (IC; log scale) of each tone-pip (from sequence onset). In REGRAN and REGxREGy, a spike in IC is seen immediately following the 1^st^ tone interrupting the regularity, and one cycle + three tones is required to discover the new regularity in REGxREGy (manifested as a drop in IC). **B:** Group-RMS of brain responses to the three sequence types. ENVnovel and ENVnovel fast responses are aligned to the time of change, marked by the black dotted line. Shaded areas around the curves represent 1 SD, computed with bootstrap resampling (1000 iterations, with replacement). Plotted data are baselined 0.1 s before stimulus onset. Statistical tests are performed on data baselined 0.1 s before the change (at 2 sec). Intervals of significant differences between conditions are marked with a line beneath the brain response, with labels indicating the specific contrasts being compared. Overall, brain response dynamics mirror the shape and timing of model predictions. **C:** Comparison of response dynamics between ENVnovel and ENVnovel fast contexts. Interruption response (REGxREGy; orange shading) minus the no-change condition (REG) in ENVnovel and ENVnovel fast (upper panel) and discovery response (REGxREGy; yellow shading) minus the random condition (REGRAN) in both contexts (bottom panel). To align the stimuli across slow and fast tone presentation rates, data are interpolated based on the number of tones presented. The black line beneath the trace indicates the intervals of significant differences during the interruption and discovery phases between ENVnovel and ENVnovel fast (in the discovery a cluster-threshold correction p < 0.1 was used instead of < 0.05 elsewhere). The black dotted lines indicate the 1^st^ and 2^nd^ cycles of REGy. **D:** MEG source localization results for ENVnovel (upper panel) and ENVnovel fast (bottom panel) in the interruption (orange shaded window) and discovery (yellow shaded window) phases identified in the time domain analyses. Group SPM t maps are shown for the significant contrasts, thresholded at p = 0.05 (uncorrected). These are superimposed on the MNI152 T1 template with the coronal and axial sections at x = 48, y = −18, and z = 3 mm.

Statistical analysis was always performed across subjects. Cluster-based permutation testing (Maris and Oostenveld, 2007) was used to account for multiple comparisons across adjacent time points within the full latency range of the epochs. Clusters of significant differences between conditions were formed by identifying adjacent time points with p-values less than 0.025. A cluster-level threshold of p < 0.05 was applied to t-statistic and the Monte Carlo method with 1000 iterations was used to estimate the null distribution of this statistic.

#### Source reconstruction

We examined the sources underlying two slow modulation phases of the sustained response upon the transition between patterns: an “*interruption*” phase which is triggered by the interruption of the initial REG, and a “*discovery*” phase which tracks the discovery of the new pattern structure following the transition. To localize the sources underlying these phases, with SPM12 (Litvak and Friston, 2008; López et al., 2014), we performed source analyses on the DSSed, low pass filtered (45Hz) signals. Sensor-level data were converted from Fieldtrip to SPM. Using 3 fiducial marker locations, the data were co-registered to a generic 8196-vertex inverse-normalized canonical mesh warped to match a T1 MRI template (Ashburner and Friston, 2005). The forward model was solved with a single shell forward head model for all subjects. Source reconstruction was performed using the multiple sparse priors (MSP) model optimized with the Greedy Search (GS) technique (Friston et al., 2008; Henson et al., 2011; López et al., 2014) by iterating over successive partitions of multiple sparse priors to find the set yielding the best fit (here we specify a total of 512 dipoles). The MSP model without Hann windowing was used to invert task-specific conditions together.

To identify areas supporting the interruption and the discovery phases, data from the ENVnovel and the ENVnovel fast contexts were averaged by condition and inverted in the interval covering two REG cycles following the interruption of the initial regular pattern (between 2 and 3 s in ENVnovel and between 1 and 1.5 in ENVnovel fast). After inversion, source estimates were averaged over the following intervals: 2.2-2.5 to cover the interruption and 2.7-3 s to cover the discovery in ENVnovel, and 1.65-1.25 to cover the interruption, and 1.39-1.5 s to cover the discovery in ENVnovel fast. These intervals were chosen to coincide with the timing of divergence between the conditions as seen in the time domain analysis (Fig. 2).

To identify areas involved in the detection of a resumed vs a novel pattern, data from REGxranREGx condition in the ENVresume context were inverted in the 0.5 to 3.5 s interval. After inversion, source estimates were averaged over a 300 ms window starting from 0.7 s to cover the detection of novel REGx, and from 2.9 s to cover the recognition of the resumed REGx after the interruption.

Source estimates were then projected to a 3D source space and smoothed [8-mm full width at half maximum (FWHM) Gaussian smoothing kernel] to create Neuroimaging Informatics Technology Initiative (NIfTI) images of source activity. These were entered into one-way within ANOVA for second-level statistical inference analysis (see Results for the specific contrasts). Statistical maps were thresholded at p < 0.05 uncorrected across the whole-brain volume.

## Results

### MEG responses reveal the dynamics of tracking changes in sequence structure

**Fig. 2A** illustrates ideal observer modeling under the ENVnovel context. When transitioning from a regular to a random pattern (REGRAN stimulus), the model output shows a sharp spike in IC (one tone after transition), driven by the violation of expectation introduced by the first tone of the RAN sequence. Subsequently, the IC stabilizes at an elevated level, reflecting the sustained unexpectedness of the RAN sequence (Fig. 2A). In contrast, transitioning from a regular to a new regular pattern initially exhibits similar dynamics to REGRAN, as the initial tones of the new REG sequence are also unexpected. However, as the new REG pattern begins repeating, the IC gradually decreases (within one cycle + three tones after the transition), reflecting growing alignment between the incoming tones and the updated context.

**Fig. 2B** (upper panel) shows MEG data for all conditions of the ENVnovel context. A series of onset responses between 0.1-0.4 s post-stimulus onset is followed by a rise to a sustained evoked response, on which responses to individual tones are superimposed. Following the interruption of REG, both REGxREGy and REGRAN conditions exhibit an abrupt drop in power at about four tones from the change (REG > REGRAN at 0.18 s, REG > REGxREGy at 0.19 s from the transition to RAN and REG respectively). This is similar to observations from a previous study (Barascud et al., 2016) and has been hypothesized to reflect the interruption, induced by the new tones, of top-down processing associated with the predictable REG pattern.

Whilst the sustained response remains low in the REGRAN condition, reflecting decreased sequence predictability compared with the regular sequence, it rises back to the REG level in the REGxREGy condition, reflecting the discovery of the new REG. REGxREGy starts diverging from REGRAN at 0.72 s from the change, indicating that it takes about one cycle + 4.4 tones to establish a new regularity model.

Overall, MEG responses exhibit similar dynamics to the ideal observer model in how they track changes in predictability. However, key differences emerge. While the model continues refining its representation of predictability—such as showing a continuous reduction of IC for REG sequences—brain responses appear to plateau after two REG cycles (approximately 1 s). Notably, the latency of REG detection following the transition aligns closely with the ideal observer’s predictions (one cycle + four tones), but the drop in the sustained response, associated with detecting the violation of regularity, lags behind the model by four tones (0.2 s). This discrepancy could reflect circuit constraints (e.g. related to the latency required to interrupt top-down processing) imposing limits on neural regularity interruption not reflected in the model. In this case, the latencies should remain fixed regardless of event presentation rate. Another possibility is that it could relate to differences in information accumulation between the model and the brain, in which case, the latency should scale with event presentation rate. These possibilities are not necessarily mutually exclusive. To test these hypotheses, we recorded responses to informationally identical stimuli in a condition where tones were shortened to 25 ms (ENVnovel fast).

### Brain response dynamics to unfolding sequences scale with tone presentation rate

In ENVnovel fast we presented the same conditions as in ENVnovel but with sequences presented at twice the speed.

In this environment (**Fig. 2B** bottom panel), compared with the no-change REG condition, REGxREGy, and REGRAN conditions exhibit a drop in power at about 0.06 s (∼ 2.5 tones) after the interruption of the initial REG (REG > REGRAN at 0.065 s, REG > REGxREGy at 0.062 s from the change). This latency is faster than that measured in ENVnovel(0.18s), demonstrating that the interruption latency is not fixed, but depends on tone presentation rate. Overall, this suggests that interrupt latency is affected by information (number of tones experienced post-interruption) and is unlikely to reflect low-level biophysical constraints.

A similar effect is observed in the discovery phase. Compared with REGRAN, in the REGxREGy, the sustained response rises back to REG level at 0.39 s from the change, about one cycle + 5.6 tones. Overall, the interruption and discovery latencies in ENVnovel fast appear to be determined by a similar amount of information as those observed in the slower sequences.

**Fig. 2C** compares the dynamics of the interrupt (top) and discovery (bottom) responses, expressed in number of tones (information), in the ENVnovel and ENVnovel fast conditions (see methods). This analysis reveals that the initial interrupt dynamics are identical between the two conditions. However, the response in ENVnovel fast exhibits a deeper trough. Similarly, for pattern discovery, there was no difference between the two conditions during the transient period (rising slope of the sustained response), suggesting similar regularity discovery dynamics. However, the sustained response after the transition to REG stabilized at a higher level in ENVnovel fast compared to the standard ENVnovel context. The increased sustained response for REG and decreased sustained response for RAN in ENVnovel fast compared to ENVnovel may stem from the highly consistent, low-SNR responses due to low inter-individual variability (Fig. S1). One possible explanation is that the 50 ms tone patterns in ENVnovel placed excessive demands on tracking mechanisms (e.g., memory, regularity detection), whereas the 25 ms patterns (with a cycle duration of 250 ms) in ENVnovel fast remained within the capacity for all subjects.

### Interruption and discovery of regularity are associated with changes in activation within a temporo-frontal-hippocampal network

#### ENV novel, interruption phase (Table 1)

**Table 1.**
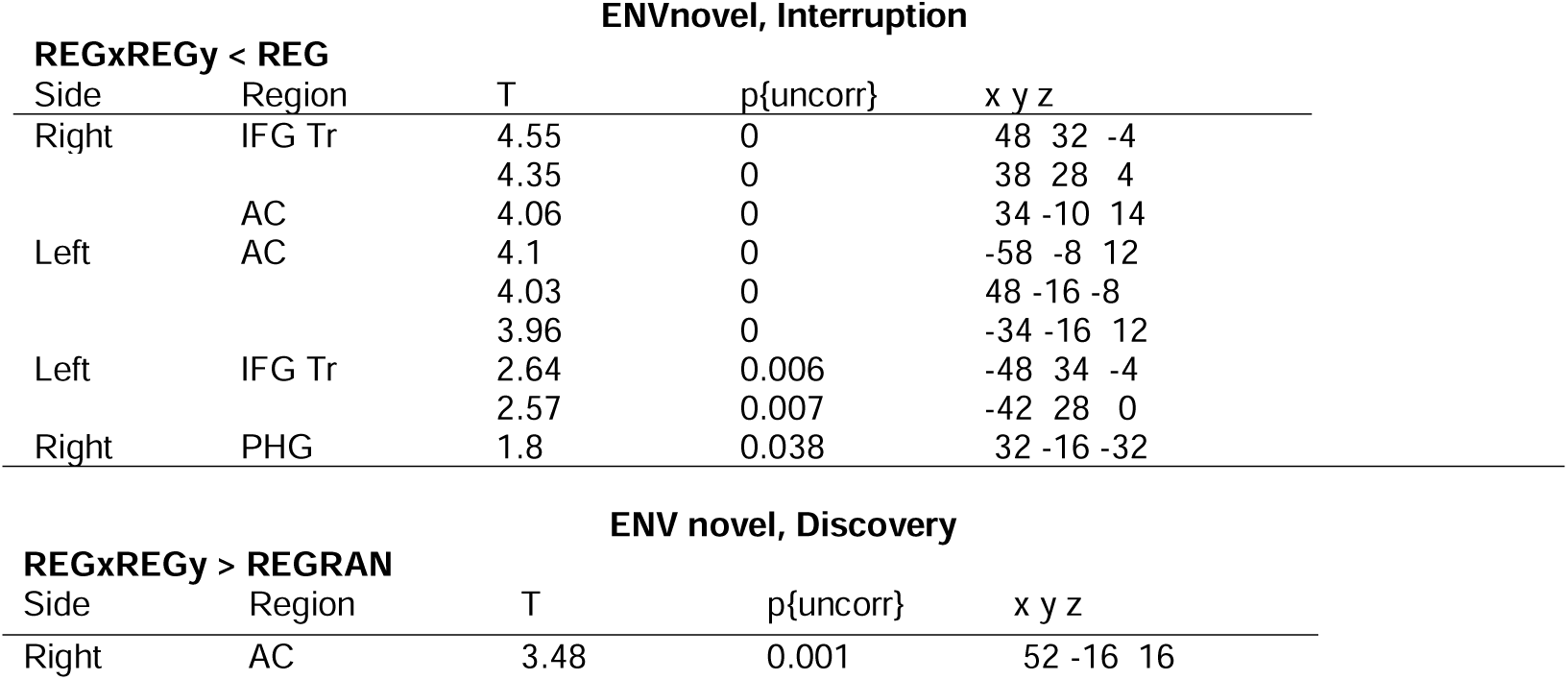
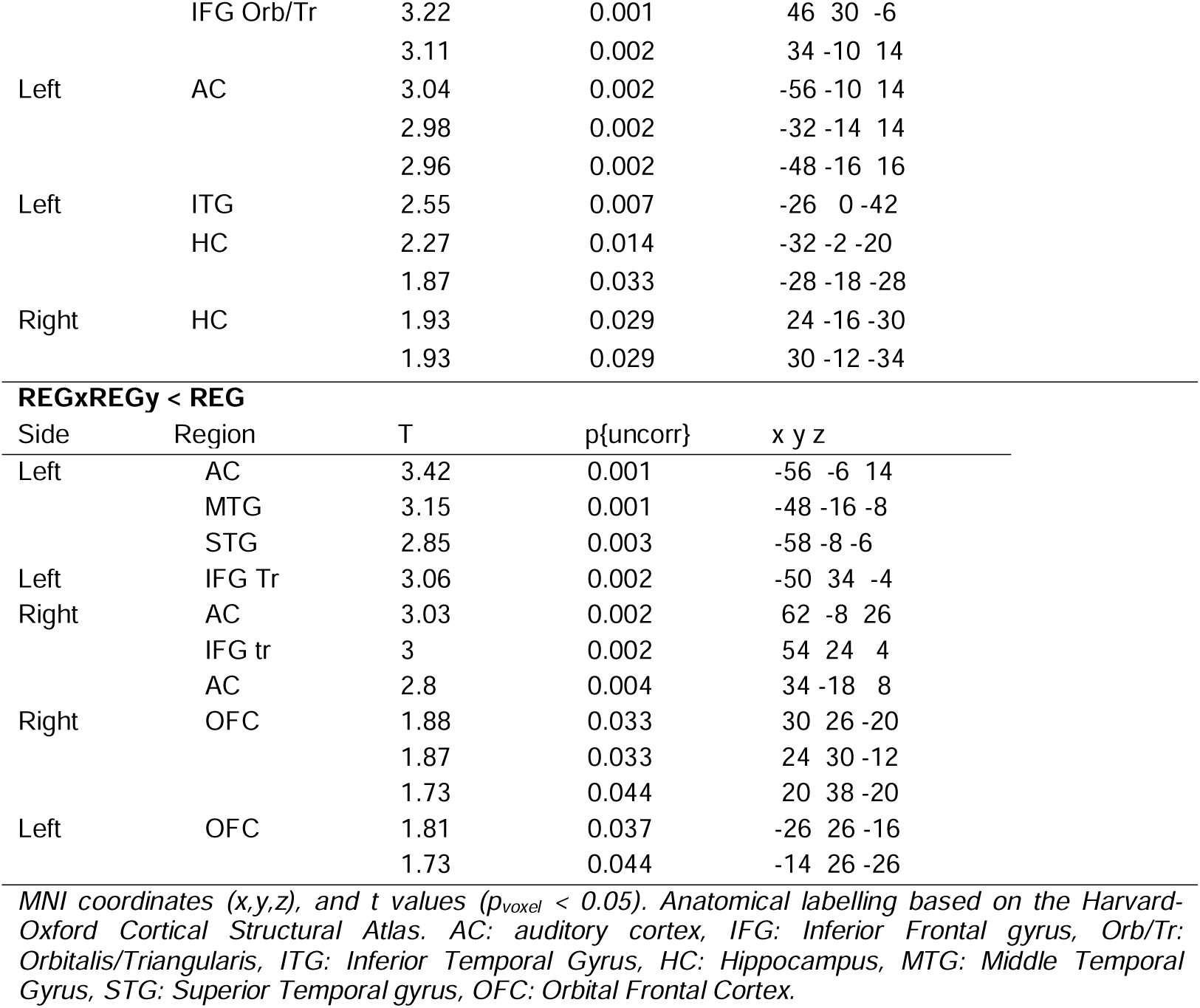
MEG source localization of REG detection and interruption in ENVnovel.

The activation map (t contrast REGxREGy < REG, p = 0.05) demonstrates reduced activity in the auditory cortex (AC, bilaterally), inferior frontal gyrus (IFG, bilaterally), and hippocampus (HC, right) following the interruption of REGx (see the pink trace in Figure 2B during the interruption phase, highlighted in orange). No areas were identified using the opposite contrast (REGxREGy > REG).

#### ENV novel, discovery phase (Table 1)

Pattern discovery is reflected in a shift in activity from a RAN to a REG state (see the pink trace in Figure 2B during the discovery phase, highlighted in yellow). This process is assessed by contrasting REGxREGy with both REGRAN and REG conditions. The activation map (t contrast, REGxREGy > REGRAN, p = 0.05; i.e. during the interval where REGy is being discovered) demonstrates increased activity in the AC (bilaterally), IFG (right), and HC (bilaterally) associated with the discovery of a new pattern. No areas were identified using the opposite contrast (REGxREGy > REGRAN). The activation map for the contrast relative to REG (t contrast, REGxREGy < REG, p = 0.05) demonstrates reduced activity in the AC (bilaterally) and IFG (bilaterally) during the discovery of REGy compared to the already established REG. No areas were identified using the opposite contrast (REGxREGy > REG).

Altogether, this pattern of source activity highlights that the presence of REG is associated with activation within a network involving temporal, hippocampal, and frontal regions. This activity fluctuates dynamically, decreasing when REG is interrupted and increasing again upon the discovery of a new REG pattern.

A similar pattern was observed when analyzing responses in the speeded (ENVnovel fast) sequences.

#### ENVnovel fast, interruption phase (Table 2)

**Table 2.**
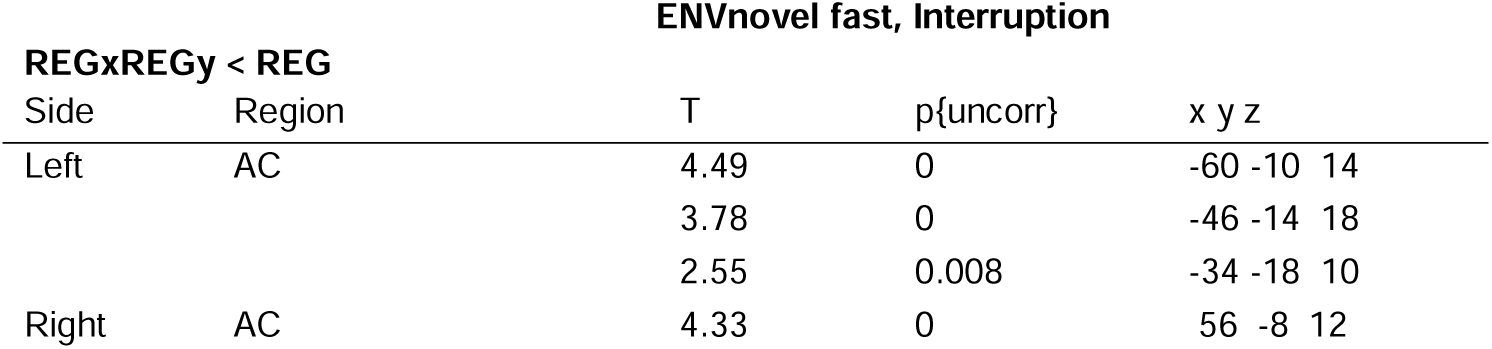
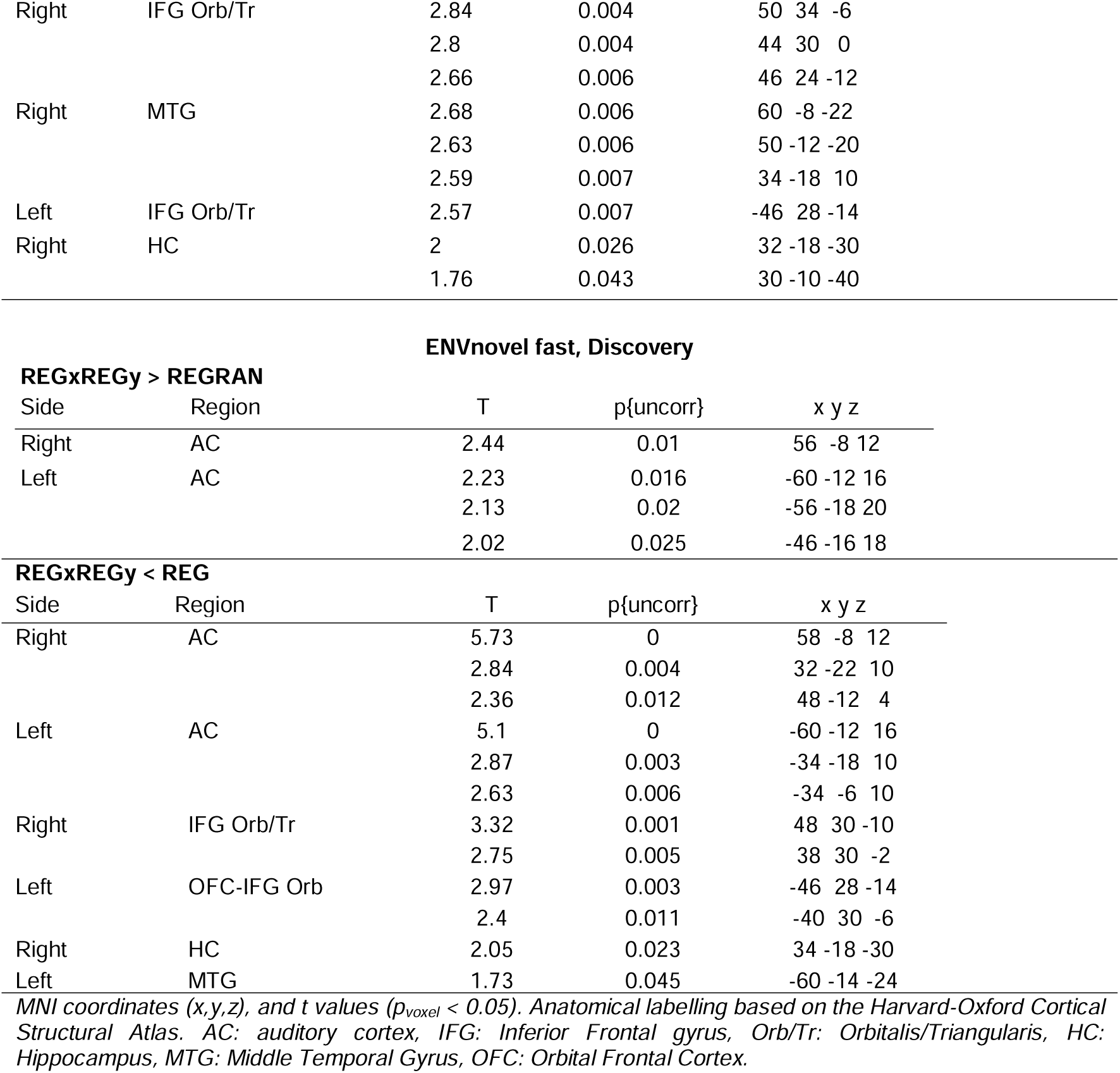
MEG source localization of REG detection and interruption in ENVnovel fast.

| ENVnovel fast, Interruption |  |  |  |  |
| --- | --- | --- | --- | --- |
| REGxREGy < REG |  |  |  |  |
| Side | Region | T | p{uncorr} | x y z |
| Left | AC | 4.49 | 0 | -60 -10 14 |
|  |  | 3.78 | 0 | -46 -14 18 |
|  |  | 2.55 | 0.008 | -34 -18 10 |
| Right | AC | 4.33 | 0 | 56 -8 12 |
| Right | IFG Orb/Tr | 2.84 | 0.004 | 50 34 -6 |
|  |  | 2.8 | 0.004 | 44 30 0 |
|  |  | 2.66 | 0.006 | 46 24 -12 |
| Right | MTG | 2.68 | 0.006 | 60 -8 -22 |
|  |  | 2.63 | 0.006 | 50 -12 -20 |
|  |  | 2.59 | 0.007 | 34 -18 10 |
| Left | IFG Orb/Tr | 2.57 | 0.007 | -46 28 -14 |
| Right | HC | 2 | 0.026 | 32 -18 -30 |
|  |  | 1.76 | 0.043 | 30 -10 -40 |

| ENVnovel fast, Discovery |  |  |  |  |
| --- | --- | --- | --- | --- |
| REGxREGy > REGRAN |  |  |  |  |
| Side | Region | T | p{uncorr} | x y z |
| Right | AC | 2.44 | 0.01 | 56 -8 12 |
| Left | AC | 2.23 | 0.016 | -60 -12 16 |
|  |  | 2.13 | 0.02 | -56 -18 20 |
|  |  | 2.02 | 0.025 | -46 -16 18 |
| REGxREGy < REG |  |  |  |  |
| Side | Region | T | p{uncorr} | x y z |
| Right | AC | 5.73 | 0 | 58 -8 12 |
|  |  | 2.84 | 0.004 | 32 -22 10 |
|  |  | 2.36 | 0.012 | 48 -12 4 |
| Left | AC | 5.1 | 0 | -60 -12 16 |
|  |  | 2.87 | 0.003 | -34 -18 10 |
|  |  | 2.63 | 0.006 | -34 -6 10 |
| Right | IFG Orb/Tr | 3.32 | 0.001 | 48 30 -10 |
|  |  | 2.75 | 0.005 | 38 30 -2 |
| Left | OFC-IFG Orb | 2.97 | 0.003 | -46 28 -14 |
|  |  | 2.4 | 0.011 | -40 30 -6 |
| Right | HC | 2.05 | 0.023 | 34 -18 -30 |
| Left | MTG | 1.73 | 0.045 | -60 -14 -24 |

MNI coordinates (x,y,z), and t values ( $p_{\text{voxel}} < 0.05$ ). Anatomical labelling based on the Harvard-Oxford Cortical Structural Atlas. AC: auditory cortex, IFG: Inferior Frontal gyrus, Orb/Tr: Orbitalis/Triangularis, HC: Hippocampus, MTG: Middle Temporal Gyrus, OFC: Orbital Frontal Cortex.

The activation map (t contrast, REGxREGy < REG, p = 0.05) demonstrates increased activity in the AC (bilaterally), IFG (bilaterally), MTG (right), HC (right) following the interruption of REGx. No areas were identified using the opposite contrast (REGxREGy > REG).

#### ENVnovel fast, discovery phase (Table 2)

The activation map (t contrast, REGxREGy > REGRAN, p = 0.05) demonstrates increased activity in the AC (bilaterally). No areas were identified using the opposite contrast (REGxREG < REGRAN) associated with the discovery of a new pattern.

The activation map (t contrast, REGxREGy < REG, p = 0.05) demonstrates decreased activity in the AC and IFG (bilaterally), as well as MTG and HC (left) during the discovery of REGy compared to the already established REG.

### Previously encountered REG patterns are detected faster than novel ones

In ENVnovel, REGx interruption was always followed by a new sequence (RAN or REGy). In ENVresume, we investigated how discovery dynamics are affected in situations where the previous REG pattern resumes after the interruption.

**Fig. 3A** shows the modeling outcomes for REGxranREGx (the original REGx pattern resumes after the interruption of 500 ms ran sequence) used in this study, compared with a hypothetical REGxranREGy condition (a new REG pattern, REGy, emerges after the interruption of 500 ms ran sequence). For the discovery of the new REGy, the model requires one cycle + four tones after the interruption to stabilize its representation. This is consistent with the estimate from Fig. 2 and the brain response timing observed for REGxREGy. In contrast, in REGxranREGx REGx is re-established far more quickly, within just three tones of the transition. This is because the model has perfect memory and retains the statistical representation of REGx in its internal store, allowing for its immediate reactivation.

**Figure 3:**
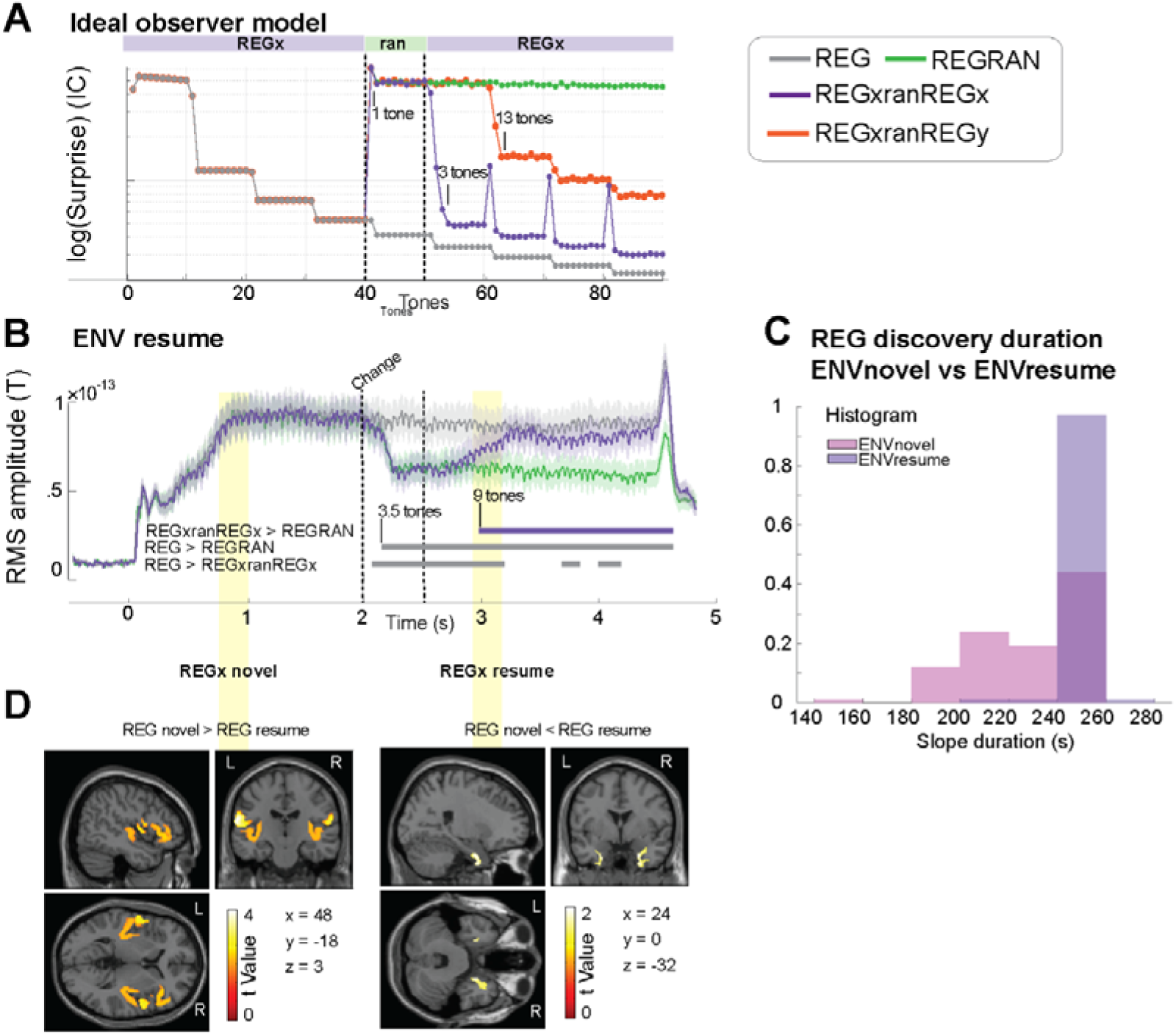
ENVresume: **A**: Ideal observer model response. The average information content (IC; log scale) of each tone-pip from sequence onset is shown for all conditions in ENVresume, together with an additional hypothetical condition, REGxranREGy (a new REG pattern, REGy, emerges after the interruption). The initial REG (REGx) is interrupted by 10 random tones (ran). The model exhibits a spike in IC following the 1^st^ RAN tone. After the interruption, it takes one cycle and three tones to discover a new REG (REGy, orange; manifested as a drop in IC) but only three tones to discover a previously present REG (REGx, purple). After REGx is resumed the model continues to exhibit “phantom” IC spikes related to the fact that the transition to ran persists in memory and affects predictability. See discussed in Magami et al (preprint) **B:** Group-RMS of brain responses. Shaded areas around the curves represent 1 SD, computed with bootstrap resampling (1000 iterations, with replacement). Plotted data are baselined 0.1 s before stimulus onset. Statistical tests are performed on data baselined 0.1 s before the change. Intervals of significant differences between conditions are marked with a line beneath the brain response trace, with labels indicating the specific contrasts being compared. **C**: Distribution of the slope duration (obtained with nested bootstrap resampling, see main text) of the discovery phase of novel REG in ENVnovel and of resumed REGx in ENVresume. **D**: Source localization results for the detection of REGx when it is novel vs when it is resumed (300 ms windows used for the localization are indicated by the yellow shadings in panel B). Group SPM t maps are shown for the significant contrasts, thresholded at p = 0.05 (uncorrected). These are superimposed on the MNI152 T1 template with the coronal and axial sections y, y, and z indicated in the figure.

**Fig. 3B** plots the MEG sustained responses. The REGxranREGx and REGRAN conditions exhibit a drop in power at about three tones after the interruption of the initial REG (REG > REGRAN at 0.18 s, REG > REGxranREGx at 0.1 s from the change). The timing of the (re)discovery of regularity in REGxranREGx was assessed with the contrast REGxranREGx > REGRAN where significant differences emerged from 0.98 s post interruption. Accounting for 500 ms of the ran sequence, the regularity has therefore been discovered in a bit less than 10 tones (i.e., before a full cycle has been completed), and about 60% faster than REGy in the ENVnovel context. The faster discovery in REGxranREGx than REGxREGy demonstrates that information previously encountered persists in memory to result in faster (by 5 tones) re-instantiation of the REG predictive model relative to a novel pattern.

The analysis above focused on the latency at which post-transition REG activation revealed a difference from the REGRAN control, that is how much evidence is accumulated before the sustained response begins to show evidence of REGx discovery -marked by the onset of the rising slope. Next, we investigated the learning rate of the REG pattern by assessing the slope of the response, namely how quickly the REGx representation stabilizes as the response reaches a plateau. The slope for the discovery of REGy in REGxREGy (ENVnovel) was quantified by subtracting the earliest significant point in the [REGxREGy > REGRAN] contrast from the latest significant point in the [REGxREGy < REG] contrast. Similarly, the slope for the (re)discovery of REGx in REGxranREGx (ENVresume) was quantified by subtracting the earliest significant point in the [REGxranREGx > REGRAN] contrast from the latest significant point in the [REGxranREGx < REG] contrast.

**Fig. 3C** plots the bootstrap-derived (100 iterations) distribution of these values (same bootstrap resampling for both environments; 1000 iterations with replacement; *p* < 0.01). We found a significant difference between the distributions (Kolmogorov-Smirnov test*, D* = 0.550, *p* < .001) and their means (Wilcoxon Signed-Rank Test, Z = -5.327, p < .001), with the mean learning duration being 0.228 ± 0.026 s in ENVnovel and 0.246 ± 0.005 s for ENVresume. This comparison revealed a shallower slope in ENVresume than ENVnovel suggesting some higher degree of memory decay for REGxranREGx than REGxREGy. We also observed a narrower distribution for REGxranREGx than REGxREGy suggesting less variability in the discovery process for the resumed REGx than for the novel REGy condition. This implies that the memory of a previously encountered REGx in the ENVresume context reduces variability in the (re)discovery process.

Overall, we observed that (a) REGx in ENVresume is discovered faster than a novel regularity (REGy) in ENVnovel (b) the rising slope associated with stabilization of REG representation was on average *shallower* but also less variable in ENVresume.

### Recognition of resumed vs novel regularity is associated with increased activation within the hippocampus

In **Fig. 3D**, we localized the sources underlying the detection of REGx in the ENVresume context when it emerges for the first time (at trial onset) vs when it is resumed following the ran interruption (time windows highlighted by yellow shadings in Fig. 3B). Activity associated with a novel REGx was assessed in the time window (between 700 and 1000 ms post sequence onset, yellow shading) where brain response to REG have been shown to begin diverging from RAN (Barascud et al., 2016; Hu et al., 2024). Activity associated with resumed REGx was assessed in a time window encompassing the rising slope after the ran interruption (between 2900 and 3200 ms post sequence onset, yellow shading). Novel REGx elicits greater activity than resumed REGx in bilateral AC and right IFG (p = 0.05). In contrast, resumed REGx vs novel REGx shows greater activity in bilateral HC (p = 0.05). (Table 3).

**Table 3.** MEG source localization of recognizing familiar vs novel REG patterns.

| REGx novel > REGx resumed |  |  |  |  |
| --- | --- | --- | --- | --- |
| Side | Region | T | p{uncorr} | x y z |
| Left | AC | 4.21 | 0 | -64 -16 18 |
|  |  | 2.5 | 0.01 | -34 -12 16 |
|  |  | 2.32 | 0.014 | -58 -10 2 |
| Right | AC | 3.61 | 0.001 | 64 -10 16 |
|  |  | 2.95 | 0.003 | 54 4 2 |
|  |  | 2.65 | 0.007 | 36 -8 16 |
| Right | IFG Tr | 2.56 | 0.008 | 44 32 -2 |
|  |  | 2.5 | 0.01 | 38 28 4 |
| REGx novel < REGx resumed |  |  |  |  |
| Left | HC | 2.29 | 0.015 | -26 2 -42 |
|  |  | 2.10 | 0.023 | -26 -2 -30 |
|  |  | 1.73 | 0.047 | -34 6 42 |
| Right | HC | 2.24 | 0.017 | 24 0 -32 |
|  |  | 2.22 | 0.018 | 24 2 -42 |

MNI coordinates (x,y,z), and t values ( $p_{\text{voxel}} < 0.05$ ). Anatomical labelling based on the Harvard-Oxford Cortical Structural Atlas. AC: auditory cortex, IFG: Inferior Frontal gyrus, Tr: Triangularis, HC: Hippocampus.

Therefore, the rediscovery of REGx is associated with reduced activity in auditory cortical and frontal sources, but increased activity in the hippocampus.

## Discussion

Sustained neural activity in response to statistical shifts in auditory sequences reflects the processing of building and updating predictive models. We examined the factors that affect this updating, revealing several key insights: (1) The timing of the sustained response drop during the *interruption phase* depends on information being accumulated, not tone presentation. This supports a “wait-and-see” strategy, where the brain holds on to existing predictive models rather than immediately discarding them. (2) Regularity discovery is faster and more consistent for previously encountered patterns, demonstrating automatic integration of both long-term and short-term statistical information for sequence structure processing. (3) Overall, the interruption and discovery dynamics remained consistent across different tone presentation rates, suggesting that these neural processes are governed by statistical structure rather than absolute timing. (4) Modulations of the sustained response consistently correspond with modulations of activity in auditory, hippocampal, and frontal regions, highlighting this network’s role in top-down information flow. Notably, the hippocampus shows greater activation when processing familiar REG patterns, reinforcing its involvement in implicit auditory memory (Billig et al., 2022).

### The sustained response closely tracks the emergence and interruption of tone patterns

MEG sustained response modulations in passive listeners mirror the pattern exhibited by an ideal observer model, but with key differences that hint at diverging heuristics: while the discovery of regularity follows timing dynamics that are nearly identical to those predicted by the model (see also Barascud et al., 2016), the latency of the interrupt response is slower than model predictions. The drop in sustained response emerges ∼200 ms (four tones) after the REG interruption. One possibility is that the delay arises from inherent circuit-level processing constraints, implying a fixed latency independent of stimulus properties (e.g., involving a neural signal potentially mediated by neurotransmission; (Yu and Dayan, 2005)) that takes ∼200 ms to develop. Alternatively, the delay may reflect a built-in “wait-and-see” period, during which the brain accumulates evidence before disengaging from a previously established model. Since abandoning a top-down expectation is risky, the brain may require sufficient evidence to confirm that the environment has changed. Under this hypothesis, the latency (and amplitude) of the interrupt response should depend on stimulus properties, such as preceding context and tone duration. Our findings support the latter hypothesis. When tone duration was reduced to 25 ms, the latency of the interrupt response decreased proportionally, suggesting that the brain employs a dynamic process to decide whether to disengage from an existing model.

As a note, in ENVnovel, REG violations always signaled a context change, while in ENVresume, the original REG resumed 50% of the time after 500ms. One might expect an observer to be sensitive to these statistics and adjust the interrupt response accordingly. However, interrupt responses remained unchanged between contexts, suggesting that global environmental regularities do not influence the response.

### Memory effects and reinstatement of previously encountered regularities

Using REGxranREGx stimuli in the ENVresume context, we investigated whether regularity discovery reflects the accumulation of local statistics or whether it draws on longer-term contextual memory. In these stimuli, REGx was interrupted by 10 random tones (500 ms) before resuming. A dependence on just local statistics would result in REGx discovery taking the same time as a novel REGy pattern. In contrast, we show, an earlier rise in the sustained response following REGx reintroduction (∼9 tones, less than one cycle) compared to novel transitions (one cycle + four tones). Previous studies have shown that familiarity speeds up brain responses to actively attended sequences (e.g., Herrmann et al., 2021). Here, we demonstrate that these effects also occur automatically in naïve, distracted listeners and during “one-shot” learning with REGx patterns that are novel on each trial. This finding suggests that the brain automatically retains a memory of the recently heard context—even when it is not behaviorally relevant—and that this memory persists despite the intervening random segment.

Compared to a perfect memory model, REGx re-discovery is slower than predicted (9-10 tones vs. 4 tones), while REGy discovery aligns with the expected onset (1 cycle + 4 tones). REGx also shows a slower learning rate than REGy, indicated by a shallower rise in the sustained response (Fig. 3C). This suggests that pattern representations in memory decay over time (Harrison et al., 2020). Accordingly, when REGy begins repeating, the brain compares the incoming information with the freshly stored REG cycle. In contrast, by the time REGx resumes after 10 random tones, its memory trace has already begun to fade.

The nature and duration of this memory store remain unclear. Previous research has demonstrated long-lasting and implicit sensory memory for reoccurring arbitrary auditory patterns (e.g., noise or tone sequences (Agus et al., 2010; Bianco et al., 2020, 2023; Agus and Pressnitzer, 2021; Herrmann et al., 2021; Ringer et al., 2022). Our findings of increased hippocampal activation to the previous vs. first encounter of REGx in REGxranREGx (see Fig. 3) suggest that this form of memory relates to the hippocampus’s role in pattern tracking (see below).

Additionally, we cannot determine whether this memory effect is specific to the ENVresume context—i.e., whether the brain learns that retaining memory is beneficial because REGx is likely to resume—or whether it occurs automatically for all unfolding sounds.

### Modeling sequence tracking

We benchmarked our hypotheses against the IDyOM model, previously shown to predict brain responses to tone patterns like those used here, as well as more natural stimuli such as music (Di Liberto et al., 2020; Kern et al., 2022; Bianco et al., 2024, 2025). In particular, we observed that model predictions regarding the latency of novel REG discovery align closely with neural responses to stimuli of varying presentation rates, from 40 Hz (ENVnovel fast) to 20 Hz (this study and Barascud et al., 2016) to 4 Hz (Hu et al., 2024), suggesting that the brain may use similar probabilistic mechanisms to infer regularities.

However, key differences emerged between the neural responses latencies we observed here and model dynamics – i.e., the brain lagged behind model predictions in detecting the REGx violation and in re-discovering the REGx after interruption. Notably, we used a specific model configuration (STM) in which memory was reset at the onset of each trial. Alternative models incorporating longer memory retention and dynamic decay might better capture the heuristics underlying human brain function.

Beyond latencies, the model also differed from brain responses in sustained IC tracking. For example, brain responses to REG patterns plateau after approximately one second, whereas model responses continue refining throughout. This difference may stem from brain response noise, a hard constraint on neural activity as represented in RMS power, or processing constraints —such as limited memory capacity—that restrict the ability to refine IC further. Notably, sustained responses differed between REG patterns of 25 ms and 50 ms tone pips, with individuals better tracking the former (Fig. S1), possibly due to better fit within relevant memory constraints.

Another instance of divergence between the model and brain responses was observed when comparing post-interruption IC and sustained responses. For example, in ENVnovel (Fig. 2), the post-interruption MEG response returned to control levels, indicating equal predictability for REGx and REGy. However, the model’s IC remained influenced by REGx memory. This carryover effect occurs because the model uses recent context for predictions, causing REGx contingencies to affect REGy. With longer context (e.g., across trials), the model’s representation stabilizes. This context-dependence is further explored in Magami et al. (pre-print), examining how the brain “rediscovers” familiar regularities.

### A fronto-temporal and hippocampal circuit underpins sustained-response modulations

Source localization demonstrates that dynamic changes in sequence predictabilities are linked to modulation of activity in the auditory cortex, inferior frontal gyrus, and hippocampus, emphasizing a distributed network responsible for encoding and dynamically tracking sequence structure.

Growing evidence indicates that the HC is essential for rapidly extracting temporal structure from the environment (Turk-Browne et al., 2010; Bornstein and Daw, 2012; Schapiro et al., 2012, 2014). Its consistent activity across multiple conditions (in this study) and previous datasets (Barascud et al., 2016; Hu et al., 2024) underscores the critical role of memory in predictive processing, where stored information guide model budling and updating. It also reinforces the HCs automatic role in tracking structure within rapidly unfolding, non-behaviorally relevant sounds (Griffiths et al., 2020; Johnson et al., 2021; Billig et al., 2025).

Furthermore, we show that the HC is more strongly activated when processing a previously encountered REGx pattern compared to its first presentation. Critically, the heightened involvement of the HC is accompanied by decreased activity in the AC and IFG. This suggests that HC recognition of the REGx pattern facilitates the reinstatement of the associated top-down model, thereby reducing resource demands in the AC and IFG.

Overall, these findings align with the HCs hypothesized role in supporting sequential structure detection via successor representation (Gershman, 2018; Ekman et al., 2023; Fang et al., 2023) and suggest its dual role in predictive coding for both encoding and re-instating previously learned regularities (Barron et al., 2020).

## Conflicts of interest

The authors declared no conflicts of interest concerning the research, authorship, and/or publication of this article.

## Acknowledgments

This work was supported by a BBSRC project grant to MC. RB is funded by the European Union (MSCA 101064334). The funders had no role in study design, data collection, and analysis, decision to publish, or preparation of the manuscript.

## Data sharing

The data reported in this manuscript alongside related information will be available on OSF upon publication.

## Supplementary material

**Figure S1.**
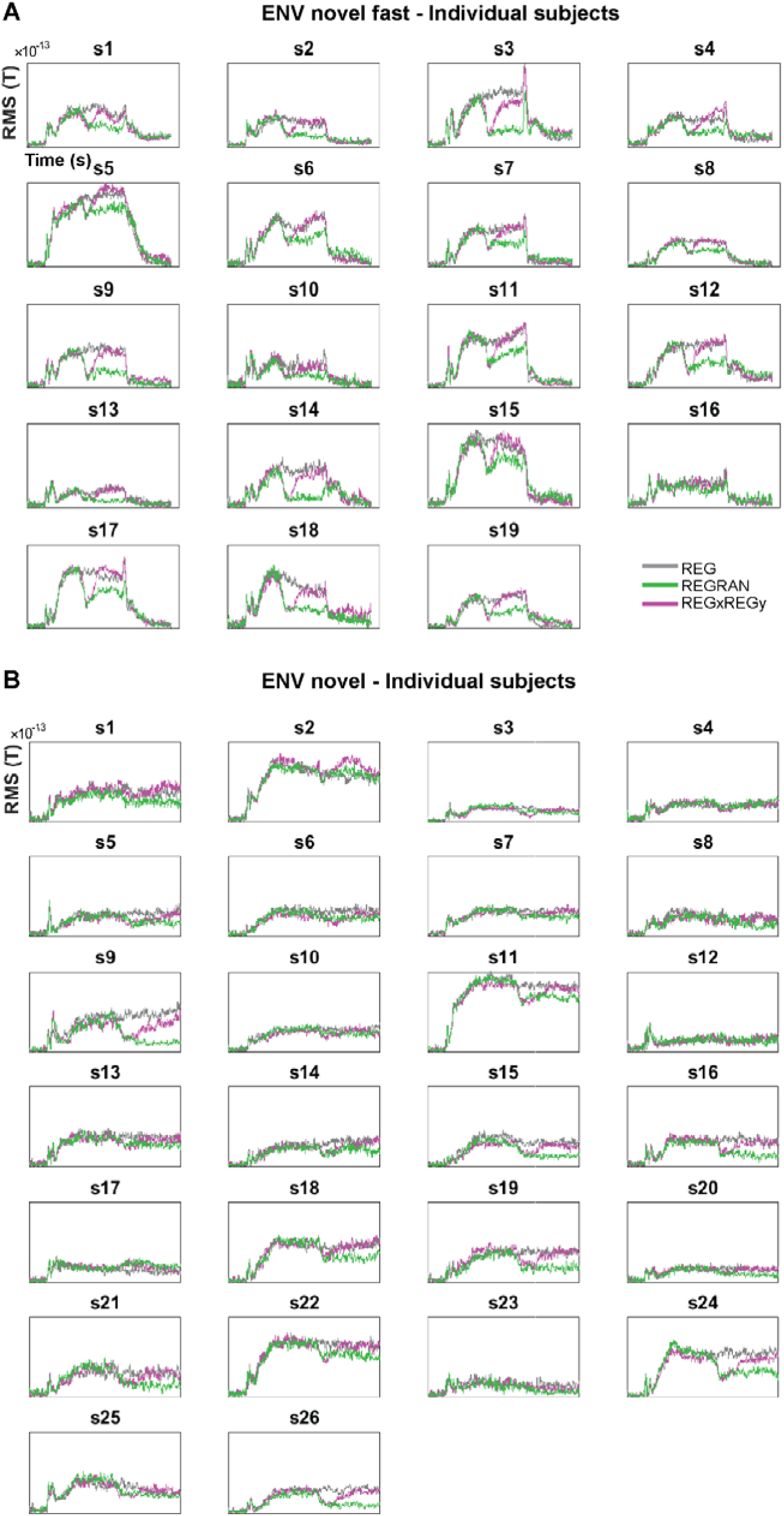
Supplementary 1. *Individual participants’ RMS of brain responses to interruption (REGxREGy and REGRAN) and discovery (REGxREGy) of a regular pattern, along with the no-change control condition (REG) in ENVnovel fast (A) and ENV novel (B) environments. Plotted data are baselined 0.1 s before stimulus onset. In ENVnovel fast, the condition difference is evident in virtually every subject, underscoring the potential of using these stimuli to understand interindividual variability in predictive and memory processes and their relationship to environmental dynamics.*

## Notes

### Competing Interest Statement

The authors have declared no competing interest.

### Summary of Updates

This version of the manuscript has been revised to update the following: - corrected typo in table 3

## References

Agus TR, Pressnitzer D (2021) Repetition detection and rapid auditory learning for stochastic tone clouds. J Acoust Soc Am 150:1735–1749.

Agus TR, Thorpe SJ, Pressnitzer D (2010) Rapid Formation of Robust Auditory Memories: Insights from Noise. Neuron 66:610–618 Available at: 10.1016/j.neuron.2010.04.014.

Al Jaja A, Grahn JA, Herrmann B, MacDonald PA (2020) The effect of aging, Parkinson’s disease, and exogenous dopamine on the neural response associated with auditory regularity processing. Neurobiol Aging 89:71–82 Available at: 10.1016/j.neurobiolaging.2020.01.002.

Ashburner J, Friston KJ (2005) Unified segmentation. Neuroimage 26:839–851.

Asokan MM, Williamson RS, Hancock KE, Polley DB (2021) Inverted central auditory hierarchies for encoding local intervals and global temporal patterns. Curr Biol 31:1762–1770.e4 Available at: 10.1016/j.cub.2021.01.076.

Barascud N, Pearce MT, Griffiths T, Friston K, Chait M (2016) Brain responses in humans reveal ideal-observer-like sensitivity to complex acoustic patterns. Proc Natl Acad Sci 113:E616–25.

Barczak A, O’Connell MN, McGinnis T, Ross D, Mowery T, Falchier A, Lakatos P (2018) Top-down, contextual entrainment of neuronal oscillations in the auditory thalamocortical circuit. Proc Natl Acad Sci U S A 115:E7605–E7614.

Barron HC, Auksztulewicz R, Friston K (2020) Prediction and memory: A predictive coding account. Prog Neurobiol 192.

Baumgarten TJ, Maniscalco B, Lee JL, Flounders MW, Abry P, He BJ (2021) Neural integration underlying naturalistic prediction flexibly adapts to varying sensory input rate. Nat Commun 12:1–14 Available at: 10.1038/s41467-021-22632-z [Accessed July 13, 2021].

Bianco R, Hall ETR, Pearce MT, Chait M (2023) Implicit auditory memory in older listeners: From encoding to 6-month retention. Curr Res Neurobiol 5:100115 Available at: 10.1016/j.crneur.2023.100115.

Bianco R, Harrison PMC, Hu M, Bolger C, Picken S, Pearce MT, Chait M (2020) Long-term implicit memory for sequential auditory patterns in humans. Elife:9:e56073.

Bianco R, Tóth B, Bigand F, Nguyen T, Sziller I, Háden GP, Winkler I, Novembre G (2025) Human newborns form musical predictions based on rhythmic but not melodic structure. bioRxiv:2025.02.19.639016 Available at: 10.1101/2025.02.19.639016 [Accessed March 20, 2025].

Bianco R, Zuk NJ, Bigand F, Quarta E, Grasso S, Arnese F, Ravignani A, Battaglia-Mayer A, Novembre G (2024) Neural encoding of musical expectations in a non-human primate. Curr Biol 34:444–450.e5.

Billig AJ, Lad M, Sedley W, Griffiths TD (2022) The hearing hippocampus. Prog Neurobiol 218:102326 Available at: 10.1016/j.pneurobio.2022.102326.

Billig AJ, Sedley W, Gander PE, Kumar S, Lad M, Mohammadi Y, Berger JI, Griffiths TD, Kingdom U, Maguire E (2025) Brain bases for navigating acoustic features. bioRxiv Prepr.

Bornstein AM, Daw ND (2012) Dissociating hippocampal and striatal contributions to sequential prediction learning. Eur J Neurosci 35:1011–1023.

Chait M, Simon JZ, Poeppel D (2004) Auditory M50 and M100 responses to broadband noise: Functional implications. Neuroreport 15:2455–2458.

de Cheveigné A (2010) Time-shift denoising source separation. J Neurosci Methods 189:113–120.

de Cheveigné A, Simon JZ (2008) Denoising based on spatial filtering. J Neurosci Methods 171:331–339.

de Lange FP, Heilbron M, Kok P (2018) How Do Expectations Shape Perception? Trends Cogn Sci 22:764–779 Available at: 10.1016/j.tics.2018.06.002.

Dehaene S, Meyniel F, Wacongne C, Wang L, Pallier C (2015) The Neural Representation of Sequences: From Transition Probabilities to Algebraic Patterns and Linguistic Trees. Neuron 88:2–19 Available at: 10.1016/j.neuron.2015.09.019.

Demarchi G, Sanchez G, Weisz N (2019) Automatic and feature-specific (anticipatory) prediction-related neural activity in the human auditory system. Nat Commun:3440 Available at: https://www.biorxiv.org/content/10.1101/266841v4.abstract?%3Fcollection=.

Di Liberto GM, Pelofi C, Bianco R, Patel P, Mehta AD, Herrero JL, de Cheveigné A, Shamma S, Mesgarani N (2020) Cortical encoding of melodic expectations in human temporal cortex. Elife:9:e51784.

Ekman M, Kusch S, de Lange FP (2023) Successor-like representation guides the prediction of future events in human visual cortex and hippocampus. Elife 12:1–19.

Fang C, Aronov D, Abbott LF, Mackevicius EL (2023) Neural learning rules for generating flexible predictions and computing the successor representation. Elife 12:1–33.

Friston K, Chu C, Mourão-Miranda J, Hulme O, Rees G, Penny W, Ashburner J (2008) Bayesian decoding of brain images. Neuroimage 39:181–205.

Friston K, Kiebel S (2009) Predictive coding under the free-energy principle. Philos Trans R Soc B Biol Sci 364:1211–1221.

Garrido MI, Kilner JM, Stephan KE, Friston KJ (2009) The mismatch negativity: a review of underlying mechanisms. Clin Neurophysiol 120:453–463 Available at: http://www.pubmedcentral.nih.gov/articlerender.fcgi?artid=2671031&tool=pmcentrez&rendertype=abstract [Accessed October 25, 2012].

Gershman SJ (2018) The successor representation: Its computational logic and neural substrates. J Neurosci 38:7193–7200.

Griffiths TD, Lad M, Kumar S, Holmes E, McMurray B, Maguire EA, Billig AJ, Sedley W (2020) How Can Hearing Loss Cause Dementia? Neuron:1–12 Available at: 10.1016/j.neuron.2020.08.003.

Harrison PMCC, Bianco R, Chait M, Pearce MT (2020) PPM-Decay: A computational model of auditory prediction with memory decay. Available at: 10.1371/journal.pcbi.1008304.

Heilbron M, Chait M (2018) Great Expectations: Is there Evidence for Predictive Coding in Auditory Cortex? Neuroscience 389:54–73 Available at: 10.1016/j.neuroscience.2017.07.061.

Henson RN, Wakeman DG, Litvak V, Friston KJ (2011) A parametric empirical bayesian framework for the EEG/MEG inverse problem: Generative models for multi-subject and multi-modal integration. Front Hum Neurosci 5:1–16.

Herrmann B, Araz K, Johnsrude IS (2021) Sustained neural activity correlates with rapid perceptual learning of auditory patterns. Neuroimage 238:118238 Available at: 10.1016/j.neuroimage.2021.118238.

Herrmann B, Johnsrude IS (2018) Neural signatures of the processing of temporal patterns in sound. J Neurosci 38:5466–5477.

Herrmann B, Maess B, Johnsrude IS (2023) Sustained responses and neural synchronization to amplitude and frequency modulation in sound change with age. Hear Res 428.

Hu M, Bianco R, Hidalgo AR, Chait M (2024) Concurrent Encoding of Sequence Predictability and Event-Evoked Prediction Error in Unfolding Auditory Patterns. J Neurosci 44:1–12.

Johnson JCSS, Marshall CR, Weil RS, Bamiou DE, Hardy CJDD, Warren JD (2021) Hearing and dementia: from ears to brain. Brain 144:391–401.

Kern P, Heilbron M, Lange FP De, Spaak E (2022) Cortical activity during naturalistic music listening reflects short-range predictions based on long-term experience. Elife:11:e80935 Available at: 10.7554/eLife.80935.

Litvak V, Friston K (2008) Electromagnetic source reconstruction for group studies. Neuroimage 42:1490–1498 Available at: 10.1016/j.neuroimage.2008.06.022.

López JD, Litvak V, Espinosa JJ, Friston K, Barnes GR (2014) Algorithmic procedures for Bayesian MEG/EEG source reconstruction in SPM. Neuroimage 84:476–487 Available at: 10.1016/j.neuroimage.2013.09.002.

Maris E, Oostenveld R (2007) Nonparametric statistical testing of EEG- and MEG-data. J Neurosci Methods 164:177–190.

Oostenveld R, Fries P, Maris E, Schoffelen JM (2011) FieldTrip: Open source software for advanced analysis of MEG, EEG, and invasive electrophysiological data. Comput Intell Neurosci 2011.

Pearce MTT (2018) Statistical learning and probabilistic prediction in music cognition: Mechanisms of stylistic enculturation. Ann N Y Acad Sci 1423:378–395.

Pezzulo G, Cisek P (2016) Navigating the Affordance LandscapeLJ: Feedback Control as a Process Model of Behavior and Cognition. Trends Cogn Sci 20:414–424 Available at: 10.1016/j.tics.2016.03.013.

Press C, Kok P, Yon D (2020) The Perceptual Prediction Paradox. Trends Cogn Sci 24:13–24.

Ringer H, Schröger E, Grimm S (2022) Perceptual Learning and Recognition of Random Acoustic Patterns. Audit Percept Cogn 00:1–23 Available at: 10.1080/25742442.2022.2082827.

Schapiro AC, Gregory E, Landau B, McCloskey M, Turk-Browne NB (2014) The necessity of the medial temporal lobe for statistical learning. J Cogn Neurosci 26:1736–1747.

Schapiro AC, Kustner L V., Turk-Browne NB (2012) Shaping of object representations in the human medial temporal lobe based on temporal regularities. Curr Biol 22:1622–1627 Available at: 10.1016/j.cub.2012.06.056.

Schröger E, Roeber U, Coy N (2023) Markov chains as a proxy for the predictive memory representations underlying mismatch negativity. Front Hum Neurosci 17:1–18.

Soltani A, Izquierdo A (2019) Adaptive learning under expected and unexpected uncertainty. Nat Rev Neurosci 20:635–644 Available at: 10.1038/s41583-019-0180-y.

Southwell R, Baumann A, Gal C, Barascud N, Friston K, Chait M (2017) Is predictability salient? A study of attentional capture by auditory patterns. Philos Trans R Soc B Biol Sci 372.

Southwell R, Chait M (2018) Enhanced deviant responses in patterned relative to random sound sequences. Cortex 109:92–103 Available at: 10.1016/j.cortex.2018.08.032.

Southwell R, Tufo C, Chait M (2024) Brain responses to predictable structure in auditory sequencesLJ: From complex regular patterns to tone repetition. :1–25.

Theunissen F, Miller JP (1995) Temporal encoding in nervous systems: A rigorous definition. J Comput Neurosci 2:149–162 Available at: https://link.springer.com/article/10.1007/BF00961885 [Accessed December 20, 2024].

Tóth B, Velősy PK, Kovács P, Háden GP, Polver S, Sziller I, Winkler I (2023) Auditory learning of recurrent tone sequences is present in the newborn’s brain. Neuroimage 281.

Turk-Browne NB, Scholl BJ, Johnson MK, Chun MM (2010) Implicit perceptual anticipation triggered by statistical learning. J Neurosci 30:11177–11187.

Wacongne C, Labyt E, Wassenhove V van, Bekinschtein T, Naccache L, Dehaene S, van Wassenhove V, Bekinschtein T, Naccache L, Dehaene S (2011) Evidence for a hierarchy of predictions and prediction errors in human cortex. Proc Natl Acad Sci 108:20754–20759 Available at: https://www.pnas.org/content/108/51/20754 [Accessed December 29, 2020].

Winkler I, Denham SL, Nelken I (2009) Modeling the auditory scene: predictive regularity representations and perceptual objects. Trends Cogn Sci 13:532–540.

Yu AJ, Dayan P (2005) Uncertainty, neuromodulation, and attention. Neuron 46:681–692.

Zhao S, Skerritt-Davis B, Elhilali M, Dick F, Chait M (2025) Sustained EEG responses to rapidly unfolding stochastic sounds reflect Bayesian inferred reliability tracking. Prog Neurobiol 244:102696 Available at: 10.1016/j.pneurobio.2024.102696.

